# Systematic Functional Characterization of Human 21st Chromosome Orthologs in *Caenorhabditis elegans*

**DOI:** 10.1101/136911

**Authors:** Sarah K. Nordquist, Sofia R. Smith, Jonathan T. Pierce

## Abstract

Individuals with Down syndrome have neurological and muscle impairments due to an additional copy of the human 21^st^ chromosome (HSA21). Only a few of ~200 HSA21 genes encoding protein have been linked to specific Down syndrome phenotypes, while the remainder are understudied. To identify poorly characterized HSA21 genes required for nervous system function, we studied behavioral phenotypes caused by loss-of-function mutations in conserved HSA21 orthologs in the nematode *Caenorhabditis elegans*. We identified ten HSA21 orthologs that are required for neuromuscular behaviors: *cle-1* (*COL18A1*), *cysl-2* (*CBS*), *dnsn-1* (*DONSON*), *eva-1* (*EVA1C*), *mtq-2* (*N6ATM1*), *ncam-1* (*NCAM2*), *pad-2* (*POFUT2*), *pdxk-1* (*PDXK*), *rnt-1* (*RUNX1*), and *unc-26* (*SYNJ1*). We also found that three of these genes are required for normal release of the neurotransmitter acetylcholine. This includes a known synaptic gene *unc-26* (*SYNJ1*), as well as uncharacterized genes *pdxk-1* (*PDXK*) and *mtq-2* (*N6ATM1*). As the first systematic functional analysis of HSA21 orthologs, this study may serve as a platform to understand genes that underlie phenotypes associated with Down syndrome.

**ARTICLE SUMMARY:** Down syndrome causes neurological and muscle dysfunction due to an extra 21^st^ chromosome. This chromosome has over 200 genes, most of which are understudied. To address this, we studied whether reducing function of these gene equivalents in the worm *C. elegans* caused neuronal or muscle defects. We identified ten genes conserved between human and worm that mediate function of behaviors. Among these, we show the uncharacterized genes *mtq-2* and *pdxk-1* are important for synaptic transmission and are exclusively expressed in nervous system. Our analysis may reveal functions of poorly studied genes that affect nervous system function in Down syndrome.

## INTRODUCTION

Down syndrome (DS) is the most common genetic cause of intellectual disability, occurring with an incidence as high as 1 in ~700 live births in the United States (Canfield et al., 2006). Although DS is defined by a variety of symptoms, all individuals with DS display varying degrees of intellectual disability, including learning and memory problems, and upon middle age they develop Alzheimer’s-like dementia (Coyle, Oster-Granite, & Reeves, 1988). In addition, all individuals with DS experience neuromuscular symptoms. For instance, DS is the leading cause of neonatal heart defects, leading to a high infant mortality rate without surgical intervention (Korenberg et al., 1994; Vis et al., 2009). Another common symptom of DS, hypotonia (muscle weakness), causes deficits in both gross and fine motor skills, leading people with DS to have trouble speaking, writing, and moving efficiently (Pitetti, Climstein, Mays, & Barrett, 1992).

For over fifty years, researchers have known that an extra copy of the 21^st^ chromosome (<u>H</u>uman <u>s</u>omatic <u>a</u>utosome <u>21</u>, HSA21) underlies DS (Jacobs, Baikie, Court Brown, & Strong, 1959). However, the precise mechanisms by which trisomy 21 causes DS-associated phenotypes are largely unknown. An early hypothesis explaining the link between DS and trisomy 21 argued that the burden of the extra genetic material strains cellular processes in the DS patient, resulting in all DS symptoms (Patterson, 2009; Shapiro, 1975). Yet research on rare individuals with partial trisomy 21 has shown that the amount of extraneous genetic material does not readily account for the collection of symptoms associated with DS nor their degree of manifestation (Korenberg et al., 1994; Lyle et al., 2009). A second hypothesis posited that a region containing about 30 protein-coding genes on HSA21, termed the <u>D</u>own <u>S</u>yndrome <u>C</u>ritical <u>R</u>egion (DSCR), is responsible for all DS-associated phenotypes. This was based on the genomic area of overlap shared between two individuals with partial trisomy 21 who still displayed many DS phenotypes (Rahmani et al., 1989). A mouse model (Ts1 Rhr) containing trisomy of only DSCR orthologous genes was developed to investigate the role of the DSCR (Olson, Richtsmeier, Leszl, & Reeves, 2004). Many, but not all of the phenotypes expected for DS were observed in this mouse model, suggesting that the DSCR is important, but not solely responsible for the whole set of DS-associated phenotypes (N. P. Belichenko et al., 2009; Olson et al., 2004). Mouse models of DS trisomic for syntenic regions of HSA21 other than the DSCR have also been instrumental in the analysis of the genetic origins of DS-related phenotypes, including craniofacial defects (Richtsmeier, Baxter, & Reeves, 2000; Richtsmeier, Zumwalt, Carlson, Epstein, & Reeves, 2002), cerebellar cell loss (Baxter, Moran, Richtsmeier, Troncoso, & Reeves, 2000), as well as synaptic and hippocampal circuit dysfunction (P. V. Belichenko et al., 2004; Demas, Nelson, Krueger, & Yarowsky, 1998; Escorihuela et al., 1995; Holtzman et al., 1996; Reeves et al., 1995).

Studies focusing on individual orthologous genes on HSA21 in animal models have also provided insight into the genetic basis of DS phenotypes. Single gene studies are attractive because the underlying genetic contributions of a phenotype are more readily dissected; single genes can also offer clearer insight into pharmacotherapeutic targeting of specific gene products (Dierssen, 2012). Determining the biological function of a single gene can be approached with either a loss-of-function or gain-of-function (e.g. overexpression) approach. Both have merit. For genes for which there is no or incomplete redundancy, loss-of-function experiments provide valuable insight into gene function. Several HSA21 orthologous genes were initially characterized in this way in invertebrate models. Early work conducted in *Drosophila melanogaster*, for instance, revealed that the gene *mnb* (*minibrain*) played a critical role in postembryonic neurogenesis (Tejedor et al., 1995). Subsequent research linked fly *mnb* to its mammalian orthologue, *DYRK1A*; and, as in fly, loss of *Dyrk1a* in mice was also shown to result in abnormal neurogenesis (Patil et al., 1995; Shindoh et al., 1996; Song, Chung, & Kurnit, 1997; Song et al., 1996). Intriguingly, overexpression of *Dyrk1a* alone in a mouse transgenic model results in neurodevelopmental delay, motor abnormalities, and spatial memory defects, which suggest a potential role in DS-associated cognitive impairment (Ahn et al., 2006; Altafaj et al., 2001). Similarly, the neuronal role of the basic helix-loop-helix protein, *SIM2* (<u>s</u>ingle-<u>m</u>inded family bHLH transcription factor <u>2</u>) on HSA21 was also initially identified in fly. Mutations in the fly ortholog, *sim*, impair development of cells in the midline of the central nervous system (Crews, Thomas, & Goodman, 1988; Thomas, Crews, & Goodman, 1988). Subsequent experiments identified the mouse homolog and found that a targeted deletion of the gene led to craniofacial malformations (H. Chen et al., 1995; Shamblott, Bugg, Lawler, & Gearhart, 2002). Overexpression of *Sim2* in mouse also recapitulated behavioral aspects of established mouse models of DS including reduced exploratory behaviors and hypersensitivity to pain (Chrast et al., 2000). Single gene studies and invertebrate models, thus, complement research with traditional DS mouse models that overexpress combinations of many HSA21 genes.

Despite this progress, the *in vivo* function of a majority of genes on HSA21 remains unknown. It is impractical to perform a systematic analysis of gene function through gene knock down in mouse models. Therefore, we set out to gain knowledge about roles of HSA21 genes by studying the function of HSA21 orthologs in the nematode *Caenorhabditis elegans*, a more tractable genetic model.

*C. elegans* is a well-established genetic model and exhibits sequence similarity with at least 42% of genes associated with human disease (Culetto & Sattelle, 2000). Additionally, the *C. elegans* nervous system is compact, with only 302 neurons (White et al., 1985). Because any one neuron may be connected to any other by only 6 or fewer synapses, defects in even a single neuron are often detectable with behavioral phenotypes such as locomotion (B. L. Chen, Hall, & Chklovskii, 2006). Proper function of the muscular feeding organ, called a pharynx, also requires precise communication between neurons and muscle. Defects in pharyngeal pumping have helped identify a conserved vesicular glutamate transporter (e.g. *eat-4*) and nicotinic cholinergic receptor subunits (Avery & You, 2012). Finally, *C. elegans* synapses are also similar to those of mammals. Genetic screens using the acetylcholinesterase inhibitor aldicarb in worm have identified genes critical for synaptic function in mammals (Richmond, 2005).

Because the *in vivo* functions of HSA21 genes have not been systematically studied, we suspected that some HSA21 genes with important neuronal or muscle roles were overlooked. The goal for our study was to determine HSA21 orthologs involved in neuronal and/ or muscle function. To this end, we analyzed behavioral phenotypes induced by loss-of-function mutations and found that disruption of several uncharacterized HSA21 orthologs in worm yielded significant neuromuscular phenotypes. To determine if any HSA21 orthologs were involved in neurotransmission, we performed a second screen on loss-of-function mutants measuring their sensitivity to the acetylcholinesterase inhibitor aldicarb. We identified three genes involved in neurotransmitter release: *unc-26 (SYNJ1)*, which has previously been characterized, and *mtq-2 (N6AMT1)* and *pdxk-1 (PDXK)*, both of which have received minimal attention. The results from our behavioral and pharmacological screens may inform predictions for which genes contribute the neuronal and muscle phenotypes associated with DS.

## MATERIAL AND METHODS

### Strains

*C. elegans* were grown on NGM (nematode growth media) agar plates seeded with OP50 bacteria at 20 °C as described (Brenner, 1974). N2 Bristol served as wild type and was also used for outcrossing (Brenner, 1974). For a list of outcrossed strains, refer to **Supplemental Table 2**. The following strains were used is this study: BB3 adr-*2(gv42)* III; CB211 *lev-1(e211)* IV; CZ18537 *zig-10(tm6127)* II; DR97 *unc-26(e345)* IV; EK228 *mbk-1(pk1389)* X; FX1756 *pad-2(tm1756)* III; FX2021 *rcan-1(tm2021)* III; FX2626 *Y105E8A.1(tm2626)* I; FX2657 *nrd-1(tm2657)* I; FX3322 *dnsn-1(tm3322)* II; FX3565 *mtq-2(tm3565)* II; FX6706 *B0024.15(tm6706)* V; FX776 *sod-1(tm776)* II; GA503 *sod-5(tm1146)* II; MT1072 *egl-4(n477)* IV; NS3026 *ikb-1(nr2027)* I; PR678 *tax-4(p678)* III; PS2627 *dgk-1(sy428)* X; RB1489 *D1037.1(ok1746)* I; RB2097 *set-29(ok2772)* I; RB2436 *cysl-4(ok3359)* V; RB2535 *cysl-2(ok3516)* II; RB870 *igcm-1(ok711)* X; RB899 *cysl-1(ok762)* X; RB979 *dip-2(ok885)* I; VC200 *rnt-1(ok351)* I; VC201 *itsn-1(ok268)* IV; VC3037 *mtq-2(ok3740)* II; VC40866 *pdxk-1(gk855208)* I; VC868 *eva-1(ok1133)* I; VC943 *cle-1(gk421)* I; VH860 *ncam-1(hd49)* X.

### Transgenic Strains

We used standard techniques to generate constructs used to make the following transgenic strains:

JPS610 *vxEx610 [mtq-2p::mtq-2::mtq-2 UTR + myo-2p::mCherry::unc-54 UTR]; mtq-2 (tm3565)*
JPS611 *vxEx611 [mtq-2p::mCherry::unc-54 UTR + unc-122p::GFP]*
JPS612 *vxEx611 [mtq-2p::mCherry::unc-54 UTR + unc-122p::GFP]; vsIs48 [unc-17::GFP]*
JPS621 *vxEx621 [mtq-2p::mtq-2::mtq-2 UTR + myo-2p::mCherry::unc-54 UTR]; mtq-2 (tm3565)*
JPS906 *vxEx611 [mtq-2p:mCherry::unc-54 UTR + unc-122p::GFP]; oxIs12 [unc-47p::GFP + lin-15(+)]*
JPS929 *vxEx929 [pdxk-1p::mCherry::unc-54 UTR]*
JPS989 *vxEx929 [pdxk-1p::mCherry::unc-54 UTR]; vsIs48 [unc-17::GFP]*
JPS990 *vxEx929 [pdxk-1p::mCherry::unc-54 UTR]; oxIs12 [unc-47p::GFP + lin-15(+)]*
JPS1044 *vxEx1044 [WRM615bF01 + myo-2p::mCherry::unc-54UTR]; B0024.15(tm6706)*
JPS1045 *vxEx1045 [WRM615bF01 + myo-2p::mCherry::unc-54UTR]; B0024.15(tm6706)*
JPS1046 *vxEx1046 [WRM061aD07 + myo-2p::mCherry::unc-54UTR]; rnt-1 (ok351)*
JPS1047 *vxEx1047 [WRM061aD07 +myo-2p::mCherry::unc-54UTR]; rnt-1 (ok351)*
JPS1048 *vxEx1048 [pdxk-1p::pdxk-1::pdxk-1UTR; myo-2p::mCherry::unc-54 UTR]; pdxk-1 (gk855208)*
JPS1049 *vxEx1049 [pdxk-1p::pdxk-1::pdxk-1UTR; myo-2p::mCherry::unc-54 UTR]; pdxk-1 (gk855208)*
JPS1051 *vxEx1051 [WRM0611dD03 + myo-2p::mCherry::unc-54UTR]; eva-1(ok1133)*
JPS1052 *vxEx1052 [WRM0611dD03 + myo-2p::mCherry::unc-54UTR]; eva-1(ok1133)*
JPS1056 *vxEx1056 [WRM069bF04 + fat-7p::GFP]; cle-1 (gk421)*
JPS1057 *vxEx1057 [WRM069bF04 + fat-7p::GFP]; cle-1 (gk421)*
JPS1058 *vxEx1058 [WRM0640bC10 + myo-2p::mCherry::unc-54UTR]; pad-2 (tm1756)*
JPS1059 *vxEx1059 [WRM0640bC10 + myo-2p::mCherry::unc-54UTR]; pad-2 (tm1756)*
JPS1060 *vxEx1060 [WRM0637bA01 + myo-2p::mCherry::unc-54 UTR]; cysl-2 (ok3516)*
JPS1061 *vxEx1061 [WRM0637bA01 + myo-2p::mCherry::unc-54 UTR]; cysl-2 (ok3516)*
JPS1062 *vxEx1062 [WRM0619dG03 + fat-7p::GFP]; ncam-1 (hd49)*
JPS1063 *vxEx1063 [WRM0619dG03 + fat-7p::GFP]; ncam-1 (hd49)*
JPS1064 *vxEx1064 [cysl-2p::mCherry::unc-54 UTR]*
JPS1065 *vxEx1065 [dnsn-1p::mCherry::unc-54 UTR]*
JPS1066 *vxEx1066 [WRM0634aG08 + myo-2p::mCherry::unc-54 UTR]; dnsn-1 (tm3322)*
JPS1067 *vxEx1066 [WRM0634aG08 + myo-2p::mCherry::unc-54 UTR]; dnsn-1 (tm3322)*

### RNA interference

We performed RNAi by feeding as described (Timmons, Court, & Fire, 2001). All RNAi clones were obtained from SourceBioScience (Nottingham, UK). First, RNAi-expressing AMP-resistant bacteria were cultured overnight at 37°C with shaking in LB broth containing ampicillin (50 mg/mL) to prevent contamination of liquid cultures. The following day, ~100 μL of bacterial liquid culture was seeded on NGM plates containing 1-mM isopropyl ß-D-1-thiogalactopyranoside (IPTG) to induce expression of exogenous RNA by the T7 promoter. Once a bacterial lawn had sufficiently grown, a mixed age population of BZ1272 *nre-1(hd20) lin-15b(hd126)* double-mutant worms was placed in a 2:1 mixture of bleach and 1-M NaOH to kill bacteria and post-embryonic worms. This strain was selected due to its heightened sensitivity to RNAi in the nervous system (Schmitz, Kinge, & Hutter, 2007). Eggs were allowed to hatch and grow over the next week at 20°C, and observed for the following week. This period covers two generations. The first generation of RNAi-treated worms experienced post-embryonic effects of the RNAi treatment, while the second generation also experienced maternal and pre-embryonic effects. We scored viability phenotypes including embryonic lethality, larval lethality, and larval arrest and sterility in either the P0 or F1 generation. Control-treated worms were fed L440 background-strain bacteria that harbored an empty RNAi vector.

### Behavioral assays

All behavioral assays were performed blind to the genotype or RNAi treatment and conducted at room temperature (~20°C).

#### Radial Dispersion Assay

To measure radial distance traveled, we placed 5-8 day-one adult worms in the center of a 10-cm diameter plate thinly seeded with OP50 bacteria. The distance traveled from the center of the plate was measured at 10 minutes (Topalidou et al., 2017). A minimum of 30 worms were tested per genotype on at least two separate days.

#### Exploration Assay

A single L4-stage worm was placed on a 3.5-cm diameter plate thinly seeded with OP50 bacteria and allowed to crawl freely. After 16 hours, we removed the worm and used the worm track to count how many of 69 squares the explored across. At least 15 worms per genotype were tested on at least two separate days.

#### Aldicarb Assay

Sensitivity to the acetylcholine esterase inhibitor aldicarb was quantified as described (Mahoney, Luo, & Nonet, 2006). At least 25 day-one adult staged worms were evaluated per trial. For the aldicarb assay using RNAi-treated worms, worms were examined at a single time point (100 minutes) on 1-mM aldicarb and scored for paralysis. Trials were performed blind to RNAi treatment and in triplicate for two generations (if viable as F1). For the aldicarb assay performed on mutant worms, the number of paralyzed worms on 1-mM aldicarb was noted every half hour for three hours. A worm was considered paralyzed when it showed neither spontaneous movement nor movement in response being prodded three times on the head and tail with a platinum wire. Assays were repeated a minimum of three times.

#### Levamisole Assay

Sensitivity to the acetylcholine receptor agonist levamisole was measured as described (Lewis, Wu, Berg, & Levine, 1980). At least 25 day-one adult staged worms were placed on plates treated with 800-μM levamisole. We scored worms for paralysis every ten minutes for one hour. Assays were performed in triplicate.

#### Pharyngeal Pumping Assay

We quantified the pumping rate of staged day-one worms by eye for thirty seconds under a stereomicroscope at x100 magnification using a handheld counter. A single pump was defined as the backward movement of the grinder (Albertson & Thomson, 1976; Raizen, Lee, & Avery, 1995). At least 30 worms per genotype were analyzed.

#### Statistical Analysis

For radial dispersion, exploration, and pharyngeal pumping assays, we compared values to the wild-type control with a one-way ANOVA with a Bonferroni correction to evaluate select comparisons or Dunnett’s test when all comparisons were to wild type. Aldicarb and levamisole responses of mutants were compared to wild-type control with a two-way ANOVA and a Bonferroni correction for select comparisons or Dunnet’s test when all comparisons were to wild type. We used GraphPad Prism7 software. To minimize false positives from our screen, we set significance to p < 0.001. For all subsequent assays, significance was set p < 0.05.

## RESULTS

### Estimate of HSA21 Protein-Coding Genes

To determine a conservative number of protein-coding genes on the human 21^st^ chromosome, we queried both Ensembl and Human Gene Nomenclature Committee (HGNC) databases and selected only those proteins reviewed by SwissProt. This public database provides manual curation and review for each protein, which ensures high quality, non-redundant entries (Consortium, 2014). This strategy yielded a total number of 213 protein-coding genes on the 21^st^ chromosome (see *Orthologs* and *Other Orthologs* sheets in **Supplemental Table 1**).

### Determining HSA21 orthologs in C. elegans

To identify putative worm orthologs of protein-coding genes on the human 21^st^ chromosome (HSA21), we relied on OrthoList, a publicly available database. OrthoList compiles results from a meta-analysis of four orthology prediction programs—Ensembl Compara, OrthoMCL, InParanoid, and Homologene (Shaye & Greenwald, 2011). OrthoList predicts at least one worm ortholog for 85 of the 213 HSA21 protein-coding genes (*blue* and *light-grey wedges* in **Figure 1A**). HSA21 encodes 48 predicted keratin proteins, which have no orthologs in worm (Shaye & Greenwald, 2011). Excluding these keratin-encoding genes, *C. elegans* has at least one predicted orthologous gene to 52% of the remaining HSA21 genes (**Figure 1A**). To focus our screen on genes that most likely share conserved functions, we chose to study orthologous genes existing in any relationship except many-to-many as defined by the InParanoid algorithm (Sonnhammer & Östlund, 2015). This included one-to-one, one-to-many, and many-to-one relationships. By this measure, InParanoid identifies 47 unique HSA21 genes with predicted orthologs in worm (three wedges highlighted by *red arc* in **Figure 1B**). These 47 HSA21 human genes are represented by 51 gene in worm.

**Figure 1:**
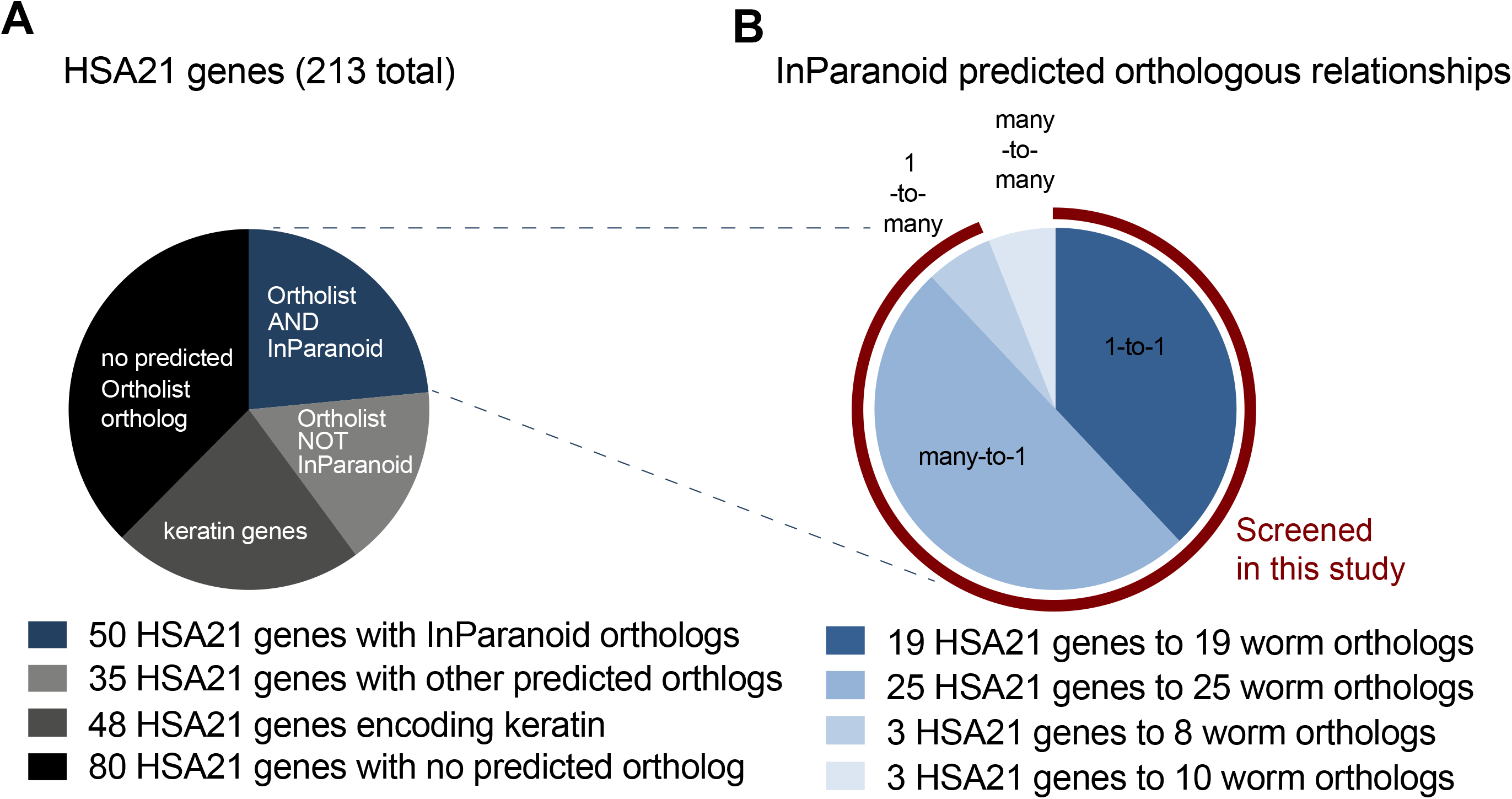
Representation of human 21^st^ chromosome genes in *C. elegans*. (**A**) HSA21 encodes a conservative estimate of 213 protein-coding genes. Excluding the 48 genes predicted to encode keratin on HSA21 (*dark gray*), over half of the remaining 165 genes have predicted orthologs in *C. elegans* as identified by OrthoList. All 85 putative orthologs in worm (*light gray* and *blue*) with an available RNAi clone were tested in the RNAi viability screen (Supplemental Table 1). (**B**) Representation of InParanoid-predicted orthologs with their human to worm relationship (*shades of blue*). We tested all predicted InParanoid orthologs except those within a many-to-many relationship with RNAi for viability and mutants for behavioral and aldicarb assays (*red*).

The imperfect evolutionary matchup of HSA21 human genes with *C. elegans* genes results in many human genes being orthologous to multiple worm genes (**Figure 1B** and see *Orthologs* sheet in **Supplemental Table 1**). For instance, the human gene *CBS* is predicted to be orthologous to four paralogs: *cysl-1, cysl-2, cysl-3*, and *cysl-4*. There was only one case of paralogous human genes in our analysis: both *RRP1* and *RRP1B* human genes are predicted to be orthologous to the single worm gene *C47E12.7*. Of the 47 HSA21 human genes, 19 are orthologous one-to-one with worm genes, 25 are orthologous many-to-one with 25 worm genes, and 3 are orthologous one-to-many with 8 worm genes.

An additional 35 HSA21 human genes had one or more orthologs predicted by the Ortholist algorithm in *C. elegans* (depicted by *light-grey wedge* in **Figure 1A** and listed in *Other Orthologs* sheet in **Supplemental Table 1**). Because most of these HSA21 human genes are predicted to be orthologous to multiple worm genes, orthology of the 35 human genes expands to represent 111 worm genes in total. For example, the human gene *ABCG1* is predicted to be orthologous to four paralogs: *wht-2, wht-3, wht-6*, and *wht-8*. In four cases, distinct but paralogous human genes are orthogous to the same worm gene. For instance, both *OLIG1* and *OLIG2* human genes are both predicted to be orthologous to *hlh-16* in worm. Due to the divergent nature of these orthologies, these 111 orthologs in worm are less likely to represent functional orthologs of HSA21 human genes. Therefore, we did not pursue them in this study.

Lastly, we found that an additional 128 protein-coding genes on HSA21 have no predicted Ortholist otholog in *C. elegans* (depicted by *black wedge* in **Figure 1A** and listed in *No Orthologs* sheet in **Supplementary Table 1**).

Taken together, our analysis above shows that discounting the keratin genes, about half of the 213 HSA21 human genes are predicted to be orthologous to genes in *C. elegans*. We focused on the set of 47 HSA21 human genes represented by 51 InParanoid orthologs in worm in this study.

### HSA21 orthologs required for viability in *C. elegans*

Before embarking on our behavioral screen using mutants, we first sought to determine which of the 51 HSA21 predicted orthologs are required for viability using RNAi (see *Orthologs* sheet in **Supplemental Table 1**). Previous large-scale and small-scale RNAi screens studied 38 of the 51 genes (Ceron et al., 2007; Cui et al., 2008; Fraser et al., 2000; Gottschalk et al., 2005; Kamath et al., 2003; Maeda, Kohara, Yamamoto, & Sugimoto, 2001; Simmer et al., 2003; Sönnichsen et al., 2005). These studies reported that RNAi treatments caused viability phenotypes for 15 of these orthologs that severely effect development, growth and/or reproduction. These include embryonic and larval lethal, larval arrest, severe growth defects, and sterility phenotypes. For an additional 24 orthologs, these studies reported that RNAi treatment caused an apparent wild-type phenotype. Out of the 51 HSA21 worm orthologs, 14 were not studied in previous RNAi screens, in part, because no RNAi reagent was available at the time.

To confirm and extend these results we performed RNAi on all 51 HSA21 orthologs for which an RNAi clone was currently available using 45 unique RNAi treatments (see *Orthologs* sheet in **Supplemental Table 1**). We found concordance for 82% or 37 out of the 45 genes tested. Only 4% or two of our RNAi treatments produced results discrepant from previous studies. RNAi treatment of *irk-1 (KCNJ6)* or *chaf-2 (CHAF1B)* were previously reported to cause severe viability phenotypes whereas we found no effect for both treatments (Gottschalk et al., 2005; Kamath et al., 2003; Nakano, Stillman, & Horvitz, 2011; Sönnichsen et al., 2005). We found that 14 out of 16 of the genes that caused inviability in our study or in previous studies when knocked down, including *irk-1* and *chaf-2*, lacked a publically available viable loss-of-function mutant. This finding is consistent with the idea that these 13 genes are probably essential. Two additional genes, *dip-2* and *dnsn-1*, had corresponding viable predicted null mutants, but we and others found that they caused inviability when knocked down by RNAi (Simmer et al., 2003; Kamath et al., 2003); perhaps this reflects a viability phenotype with low penetrance. We acknowledge that additional genes may later prove to represent essential genes too, especially if the gene lacks a corresponding null mutant, e.g. *pdxk-1*. Thus, overall our RNAi screen confirms most viability phenotype results from previous studies and suggests that at least 33% or 15 out of 45 of tested HSA21predictyed orthologs are inviable when reduced in function in *C. elegans*.

### HSA21 orthologs required for neuromuscular behaviors

Having identified which HSA21 orthologs are probably essential, we turned next to investigate the *in vivo* function of nonessential HSA21 orthologs. We tested every publically available, viable mutant, which represented 27 genes in total (see *Orthologs* sheet in **Supplemental Table 1**). We performed a phenotypic screen, selecting a battery of three behavioral assays to probe neuromuscular function. Overall, we identified ten mutants that were significantly defective in performance in these assays. To better link genotype to phenotype, we generated transgenic rescue lines for any mutant deficient in any of the three behaviors and/or tested additional mutant alleles if available.

We first examined short-term locomotion with a radial dispersion assay, which provides a general metric of locomotor function over 10 minutes. In this assay, the distance displaced over time reflects a combination of locomotor traits including frequency of body bends, reversal rate, and efficient coordination (Topalidou et al., 2017). Aside from the well-characterized uncoordinated mutant *unc-26*, an ortholog of synaptojanin (Harris, Hartwieg, Horvitz, & Jorgensen, 2000), we also identified two other mutants with rescuable deficits in radial dispersion: *cysl-2 (CBS)* and *rnt-1 (RUNX1)* (**Figure 2**).

**Figure 2:**
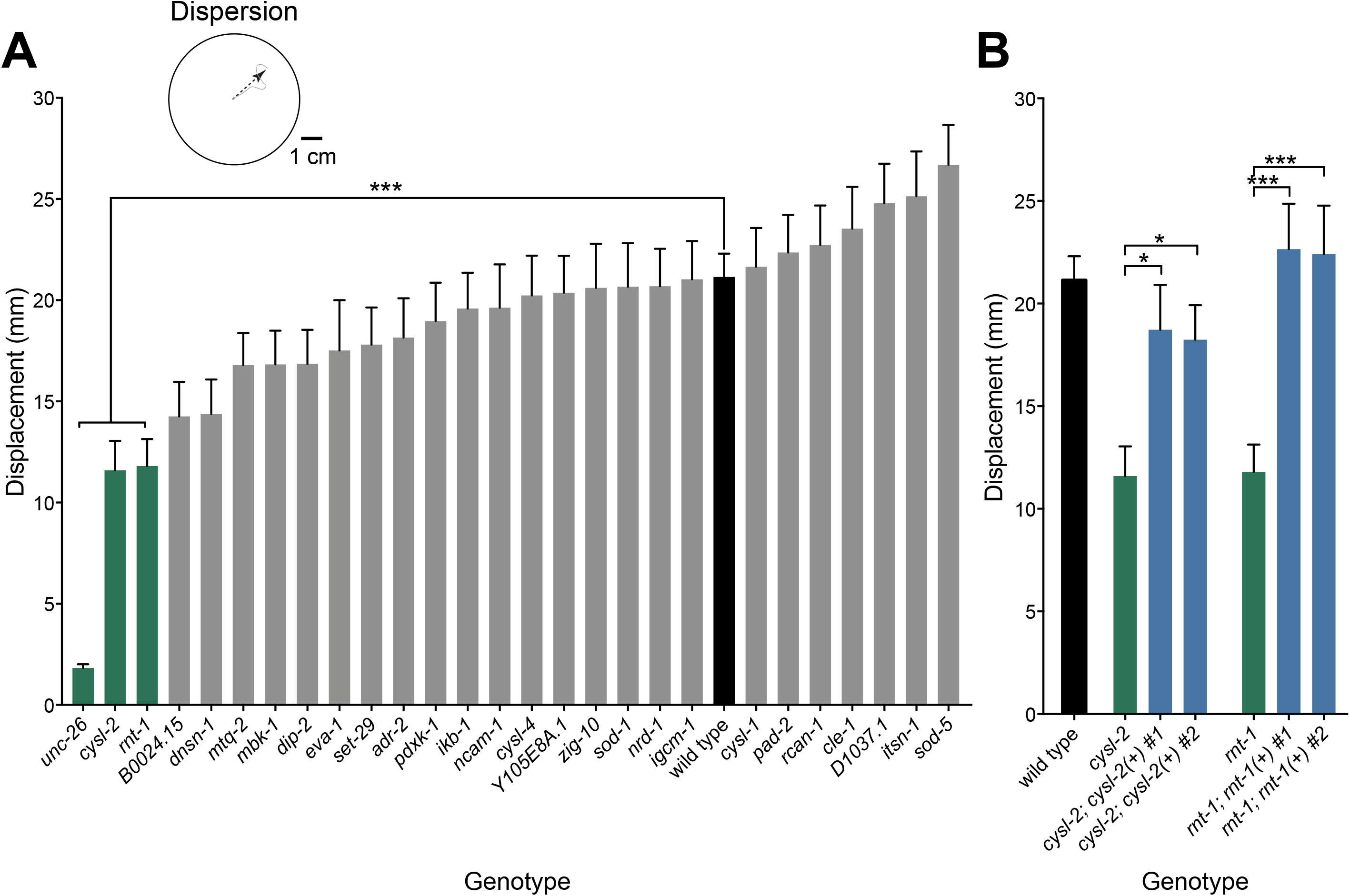
Screen for radial dispersion defects. Histograms show the mean displacement +/- SEM from start over ten minutes for each worm during a radial dispersion assay. (**A**) Three mutants (*green*) showed substantial reductions in dispersal relative to wild type (*black*). ***p < 0.001, n > 30. (**B**) Reductions in dispersal were rescued for the *cysl-2* and *mt-1* mutants (*green*) by extrachromosomal expression of the wild-type genes (*blue*). Wild-type performance shown for comparison (*black*). *p < 0.05, **p <0.01, ***p <0.001, n > 30.

Second, we examined the longer-term behavior of mutants with an exploration assay. This assay measures not only the general locomotor ability of worms, but also their tendency to explore a cultivation plate seeded with bacterial food over the course of 16 hours. While foraging, worms alternate between two states: roaming—an active state characterized by a burst of locomotion associated with a search for food—and dwelling—a more passive state associated with feeding or resting after satiation. Time spent in roaming and dwelling states depends on integrating internal neuromodulatory cues with external sensory cues. The absence of such sensory transduction leads to extended dwelling as observed in the *tax-4* mutant, which encodes a cyclic nucleotide-gated channel subunit (Fujiwara, Sengupta, & McIntire, 2002; Greene et al., 2016). In contrast, mutants with constitutive sensory input, such as the *egl-4* mutant, show extended roaming (Fujiwara et al., 2002) Extended dwelling or roaming compared to wild-type behavior can be quantified by counting the number of squares that a worm traverses over 16 hours (**Figure 3**).

**Figure 3:**
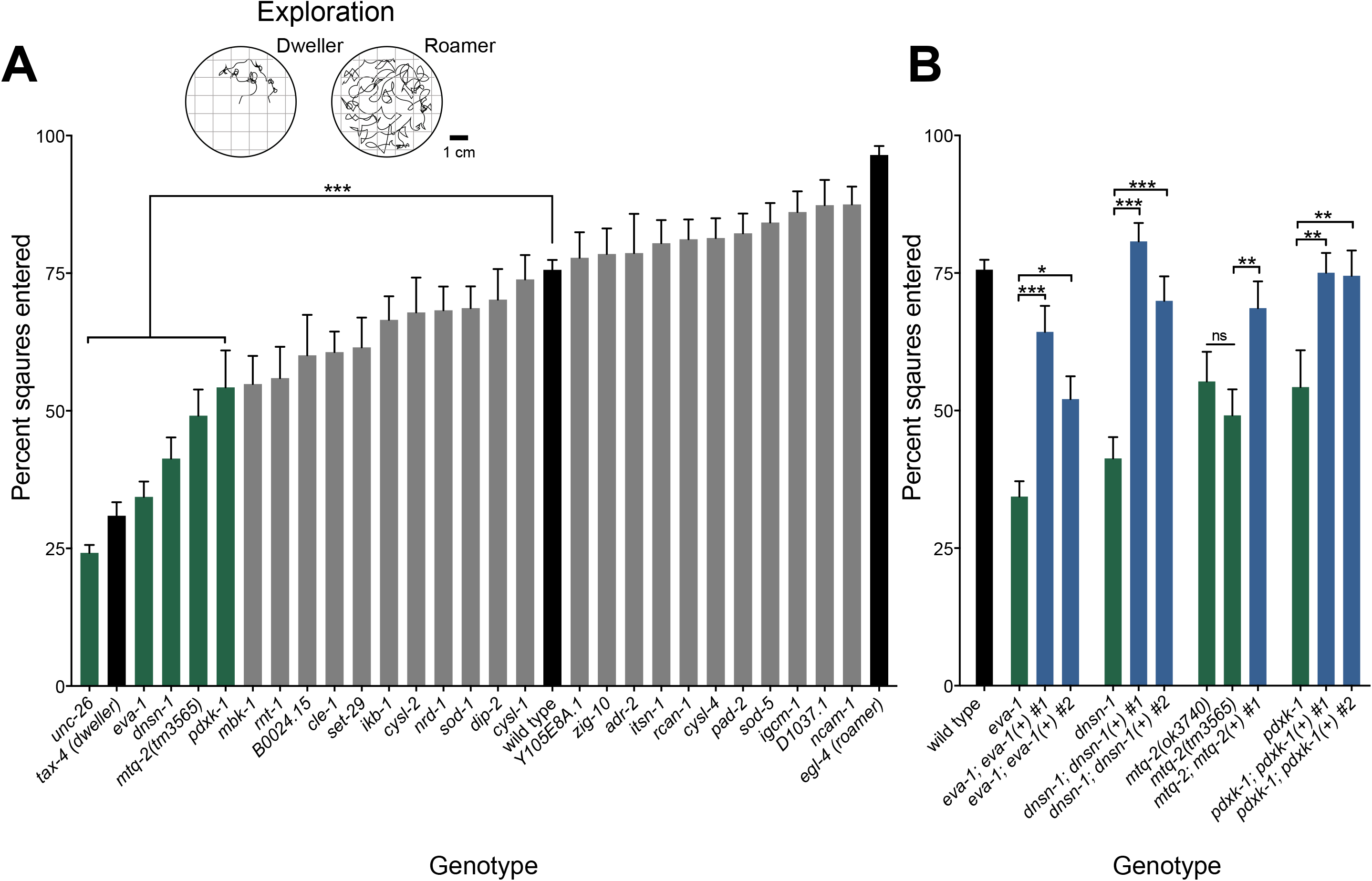
Screen for exploration defects. Histograms show the mean percentage of the assay plate traversed (percent of squares entered +/- SEM) by individual worms over a 16-h exploration assay. (**A**) Five mutants (*green*) covered significantly less area relative than wild type (*black*). ***p < 0.001, n > 15. (**B**) Reductions in exploration were rescued in the *eva-1, C24H12.5, mtq-2(tm3565)*, and *pdxk-1* mutants (*green*) by extrachromosomal expression of each of the wild-type genes (*blue*). Wild-type performance shown for comparison (*black*). *p < 0.05, **p < 0.01, ***p < 0.001, n >15.

As expected, we found that the uncoordinated mutant *unc-26 (SYNJ)* moved poorly in the exploration assay. We also identified the following four mutants with rescuable defects in extended dwelling: *eva-1 (EVA1C), dnsn-1 (DONSON), mtq-2 (N6AMT1)*, and *pdxk-1 (PDXK1)* (**Figure 3A**). These mutants appeared to move superficially similar to wild type, suggesting sensory-motor integration deficits. We rescued the exploration behavior of each mutant by transformation with wild-type copies of corresponding transgenes (**Figure 3B**).

To obtain a measure of neuromuscular activity that was distinct from locomotion, we performed a feeding assay. *C. elegans* pumps bacterial food into its gut through a feeding organ called a pharynx that uses muscles and neurons distinct from than those needed for locomotion. The rhythmic contractions and relaxations of the pharynx require precise coordination between pharyngeal neurons and muscle. We identified the following four genes required for normal pharyngeal pumping: *cle-1 (COL18A1), mtq-2 (N6AMT1), ncam-1 (NCAM2)*, and *pad-2 (POFUT2)* (**Figure 4A**). Independent null alleles of *mtq-2* displayed similar deficits in pharyngeal pumping (**Figure 4B**). Transformation with wild-type copies of each gene rescued the slow pumping phenotype of each mutant (**Figure 4B**).

**Figure 4:**
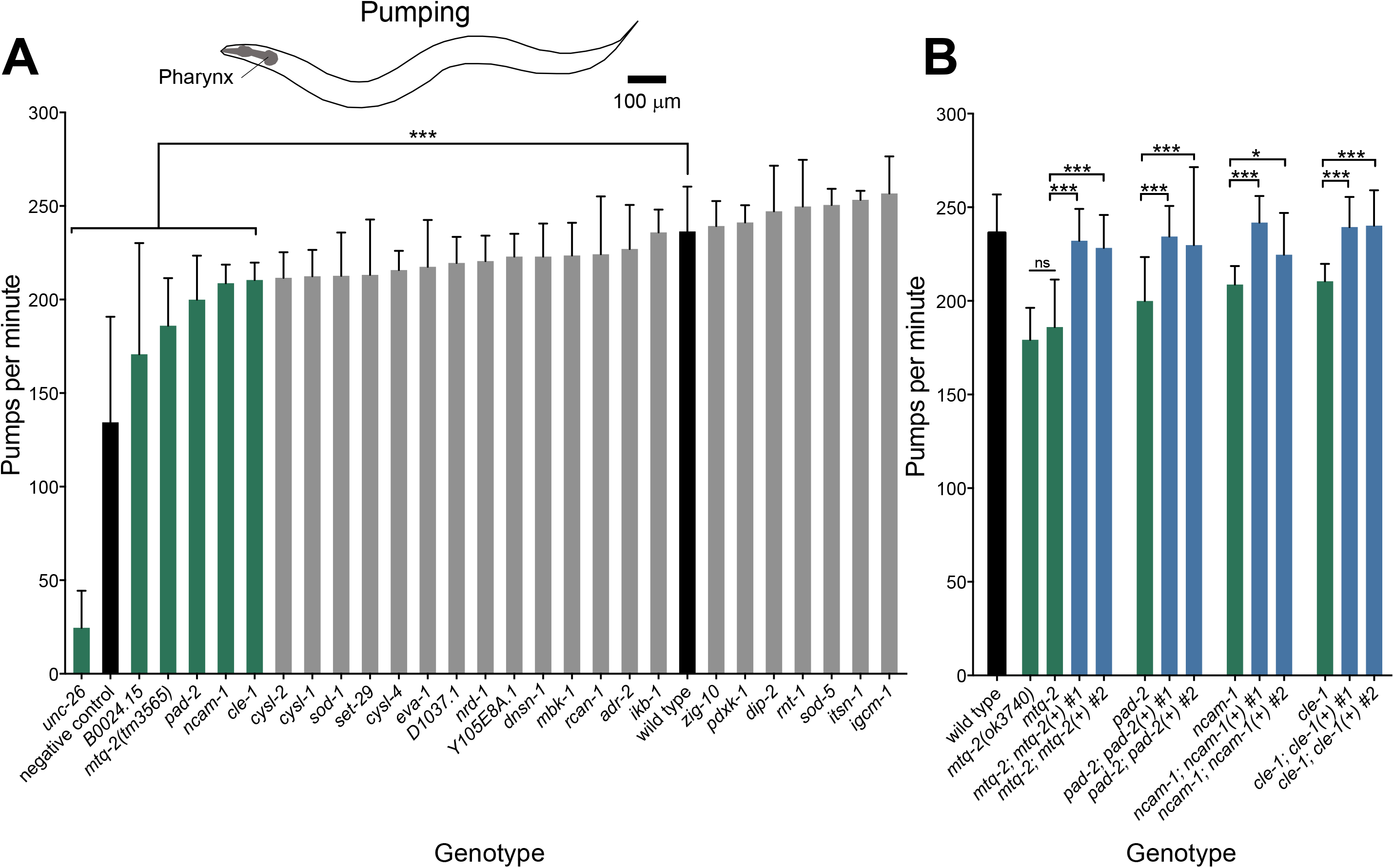
Screen for pumping defects. Histograms showing mean pharyngeal pumps per minute (+/- STD) in the presence of standard bacterial food. (**A**) Seven mutants (*green*) showed significant defects in pharyngeal pumping relative to wild type (*black*). ***p< 0.001, n > 30. (B) Mutants with pharyngeal pumping defects *mtq-2(tm3565), pad-2, ncam-1*, and *cle-1* (green) were rescued by extrachromosomal expression of the wild-type genes (*blue*). We did not observe rescue of *B0024.15*. *p < 0.05, **p < 0.01, ***p < 0.001, n > 30.

In summary, by screening three behaviors, we identified ten genes total with rescuable defects in one or more behaviors designed to probe neuromuscular function. Several among these genes—*dnsn-1 (DONSON), pdxk-1 (PDXK)*, and *mtq-2 (N6AMT1)*—are poorly characterized, yet may mediate nervous system function, development, or both.

### HSA21 orthologs required for proper synaptic function

To determine the potential role of HSA21 predicted orthologs in neurotransmission, we tested whether the same set of loss-of-function mutants displayed altered sensitivity to paralysis by the acetylcholinesterase inhibitor aldicarb. Many genes critical for key steps in synaptic function have been identified by testing mutants for altered sensitivity to aldicarb (Richmond, 2005). Aldicarb causes paralysis of worms by chronically activating body-wall muscle after prolonged presence of acetylcholine at the neuromuscular junction. Mutants defective in any aspect of synaptic transmission show altered sensitivity to paralysis by aldicarb. For instance, mutants defective in acetylcholine release show resistance to paralysis by aldicarb due to decreased levels of acetylcholine in the cleft (Mahoney et al., 2006). Conversely, mutations that increase acetylcholine release lead to increased sensitivity to paralysis (Gracheva et al., 2006; McEwen, Madison, Dybbs, & Kaplan, 2006).

We identified three mutants that displayed significant resistance to paralysis by aldicarb: *unc-26 (SYNJ1), mtq-2 (N6AMT1)*, and *pdxk-1 (PDXK)*. Compared to wild type, these three mutant strains took longer to paralyze when treated with aldicarb (**Figure 5A** and **Figure 6**). By contrast, 24 other mutants became paralyzed at a similar rate as wild-type worms. Although we measured the 180-minute time course of paralysis for all 27 mutants, we plot only the percent moving on aldicarb at the 180-minute time point for ease of comparison (**Figure 5B**). Transformation of *mtq-2* and *pdxk-1* mutants with wild-type copies of these genes rescued aldicarb sensitivity for each mutant (**Figure 6**). As a complementary strategy to the mutant screen, we also tested aldicarb sensitivity for HSA21 predicted orthologs using RNA interference. With this independent approach, we confirmed the aldicarb resistance of *pdxk-1 (PDXK-1)* and observed suggestive results for *mtq-2 (MTQ-2)* (**Supplemental Graph 1**).

**Figure 5:**
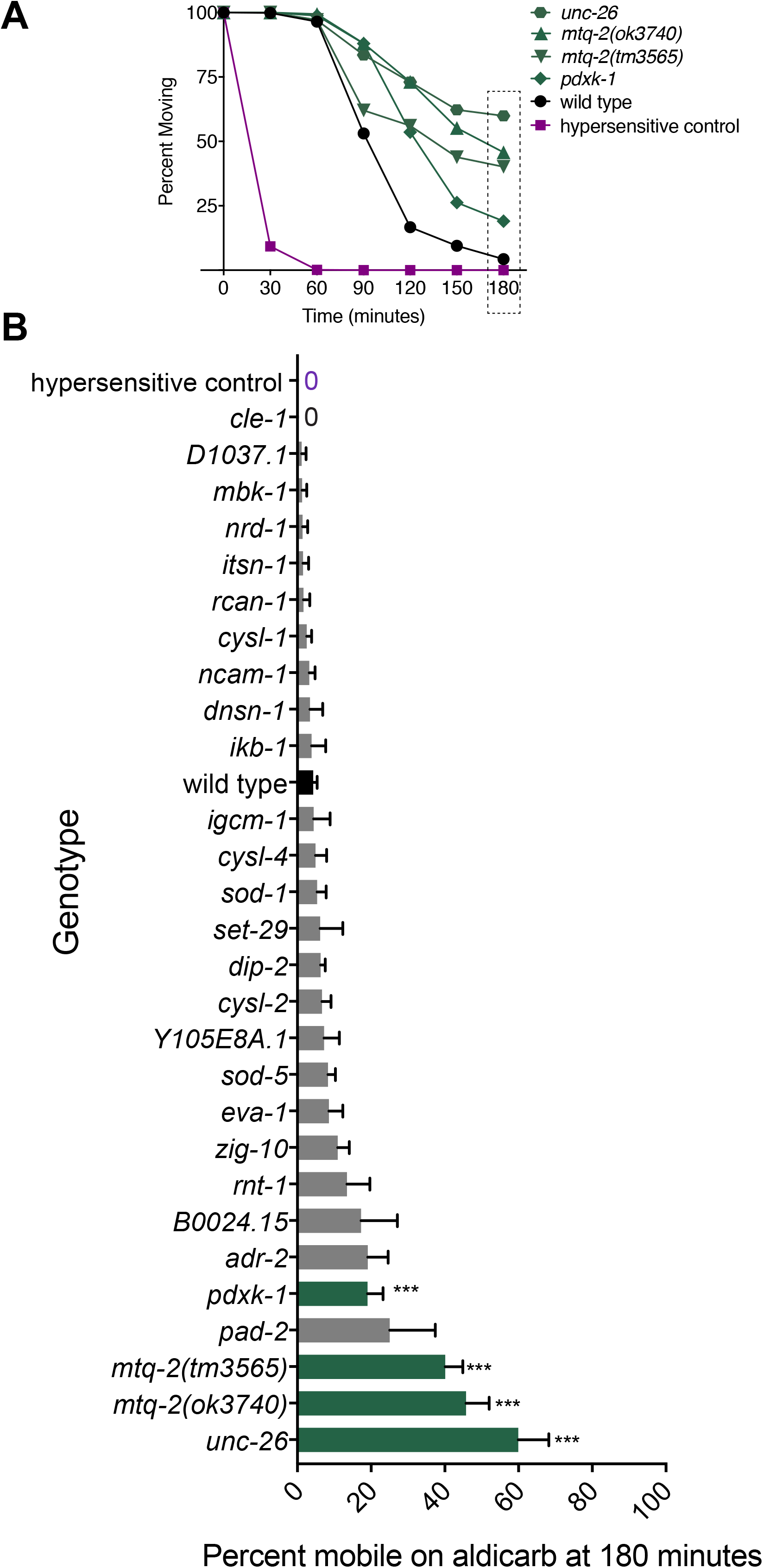
Screen for aldicarb resistance. We measured the full 180-minute time course to paralysis for 28 strains including wild type and hypersensitive control (*dgk-1*) in 1-mM aldicarb. Significant difference in paralysis rate was determined from WT with a two-way ANOVA using Dunnett’s correction for multiple comparisons. No mutants showed significant hypersensitivity. (**A**) Mean percent (+/- SEM) of animals moving on aldicarb over minutes for WT, positive control, and mutants that were significantly different from WT: *pdxk-1(gk855208), mtq-2(tm3565), mtq-2(ok3740)*, and *unc-26(e345)*. p < 0.001, n > 3. (B) Mean percent of (+/- SEM) of animals moving on aldicarb at the 180 minute time point for all mutants tested including those (*green*) that were significantly different from WT (*black*). Zero indicates that no animals were moving.

We found that RNAi treatment targeting *ncam-1* provided resistance to aldicarb; however, we also found that a null allele of *ncam-1* failed to show any resistance to paralysis by aldicarb (**Figure 4B** and **Figure 5**) suggesting that our RNAi result for *ncam-1* was a false positive. This result is consistent with a previous RNAi study that found negative results for *ncam-1* on aldicarb (Sieburth et al., 2005).

Several decades of studies have concluded that resistance to aldicarb is primarily caused by presynaptic deficiencies in acetylcholine release or postsynaptic deficiencies in receptor response (Mahoney et al., 2006). To distinguish between these possibilities, we measured the sensitivity of mutants to paralysis by levamisole, a potent agonist for the nicotinic cholinergic receptor (Lewis et al., 1980). Levamisole causes paralysis of wild-type worms by chronically activating body-wall muscle irrespective of synaptic release defects. Mutants resistant to paralysis by levamisole are defective in cholinergic receptor signaling or muscle function.

We found that *mtq-2, unc-26*, and *pdxk-1* mutants all displayed wild-type-like or hypersensitive responses to levamisole (**Figure 6**). These results suggest that resistance to aldicarb among these three mutants is due to defective presynaptic release of neurotransmitter.

**Figure 6.**
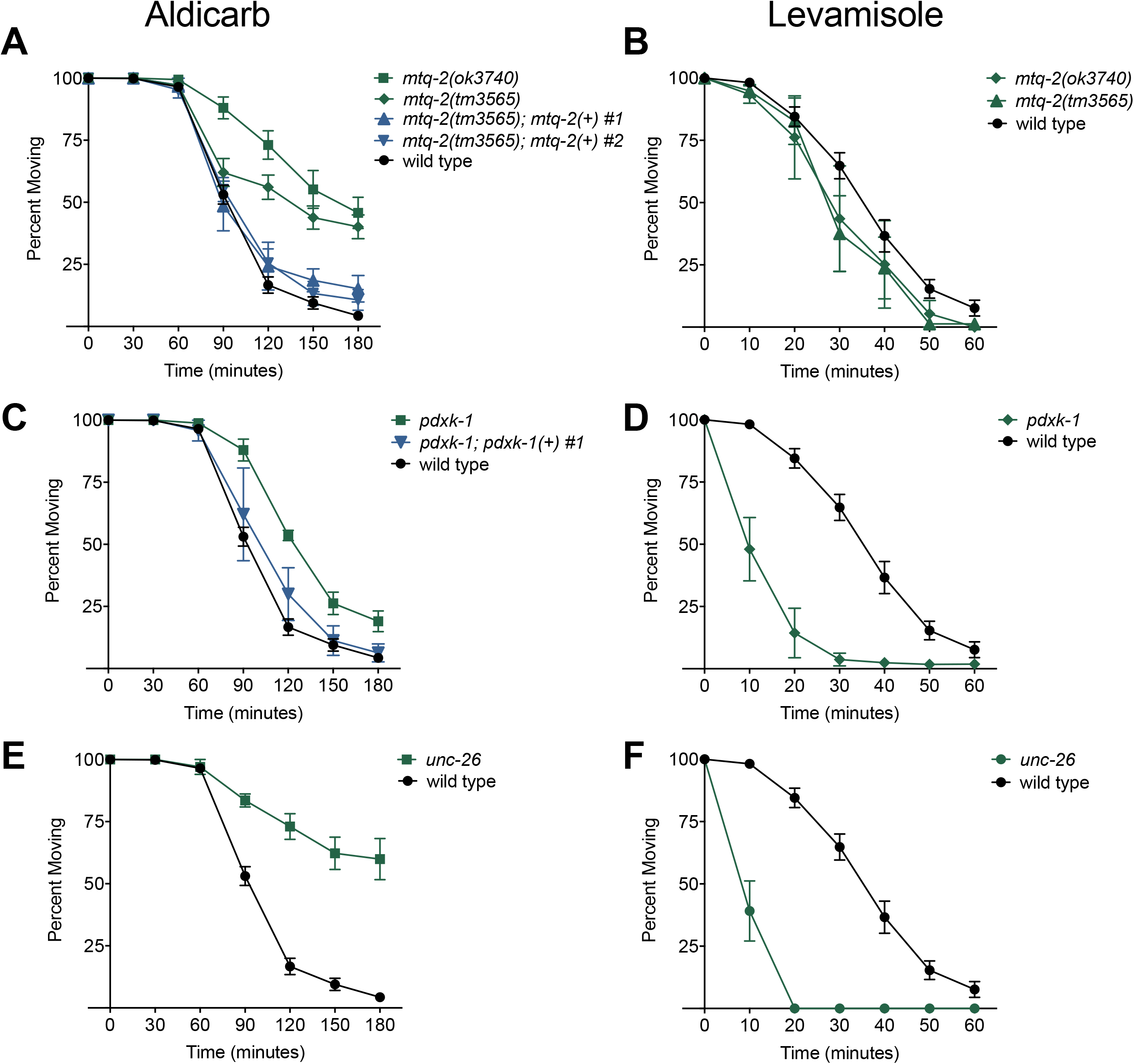
Response of deletion and putative loss-of-function mutants to aldicarb and levamisole. Time course to paralysis on 1-mM aldicarb or 800μM levamisole for aldicarb resistant mutants identified in screen (n > 3 trials, ~25 worms per trial). Wild-type controls were assayed in parallel. (**A**) Two independent alleles of *mtq-2* (*green*) displayed significant resistance to aldicarb, p < 0.001. Two of two rescue lines of *mtq-2(tm3565)* (*blue*) displayed significantly sensitivity from the mutant background, p < 0.05. (**B**) Two independent alleles of *mtq-2* displayed sensitivity to levamisole. (**C**). *pdxk-1(gk855208)* displayed significant resistance to aldicarb, p < 0.001. One of two rescue lines of *pdxk-1(gk855208)* (blue) was significantly improved from the mutant background, p < 0.01. (**D**) *pdxk-1(gk855208)* displayed hypersensitivity to levamisole relative to wild type, p < 0.001. (**E**) *unc-26(e345)* displayed significant resistance to aldicarb, p < 0.001 and (**F**) hypersensitivity to levamisole, p < 0.001.

### Expression pattern of HSA21 orthologs

We identified nine genes linked to rescuable neuromuscular defects in our study (**Table 1**). Five genes were previously shown to be expressed in the nervous system: *cle-1 (COL18A1), eva-1 (EVA1C), ncam-1 (NCAM2), pad-2 (POFUT2), rnt-1* (RUNX1) and *unc-26 (SYNJ)* (Ackley et al., 2001; Fujisawa, Wrana, & Culotti, 2007; Hunt-Newbury et al., 2007; McKay et al., 2003; Menzel et al., 2004; Schwarz, Pan, Voltmer-Irsch, & Hutter, 2009). An additional four genes had no reported expression patterns: *cysl-2 (CBS), dnsn-1 (DONSON), mtq-2 (N6AMT1)*, and *pdxk-1 (PDXK)*. To identify the tissues in which these genes may function, we generated mCherry transcriptional reporter strains. We found that *mtq-2* expressed exclusively throughout the nervous system. Expression overlapped with a subset of cholinergic neurons in the head and throughout the ventral nerve cord as defined by the *Punc-17::GFP* reporter (**Figure 7A4**). Expression of *mtq-2* mCherry reporter was not observed in GABAergic neurons defined by the *Punc-47:GFP* reporter (**Figure 7A5**). Interestingly, our mCherry reporter for *pdxk-1* expressed in GABAergic, but not cholinergic neurons (**Figure 7B**). Our transcriptional reporter for *dnsn-1* expressed robustly within the developing embryo, but not within the adult nervous system (**Figure 7C**). This suggests a potential developmental role for this gene. Lastly, we found that our mCherry *cysl-2* reporter expressed broadly throughout head and body-wall muscle (**Figure 7D**).

**Figure 7.**
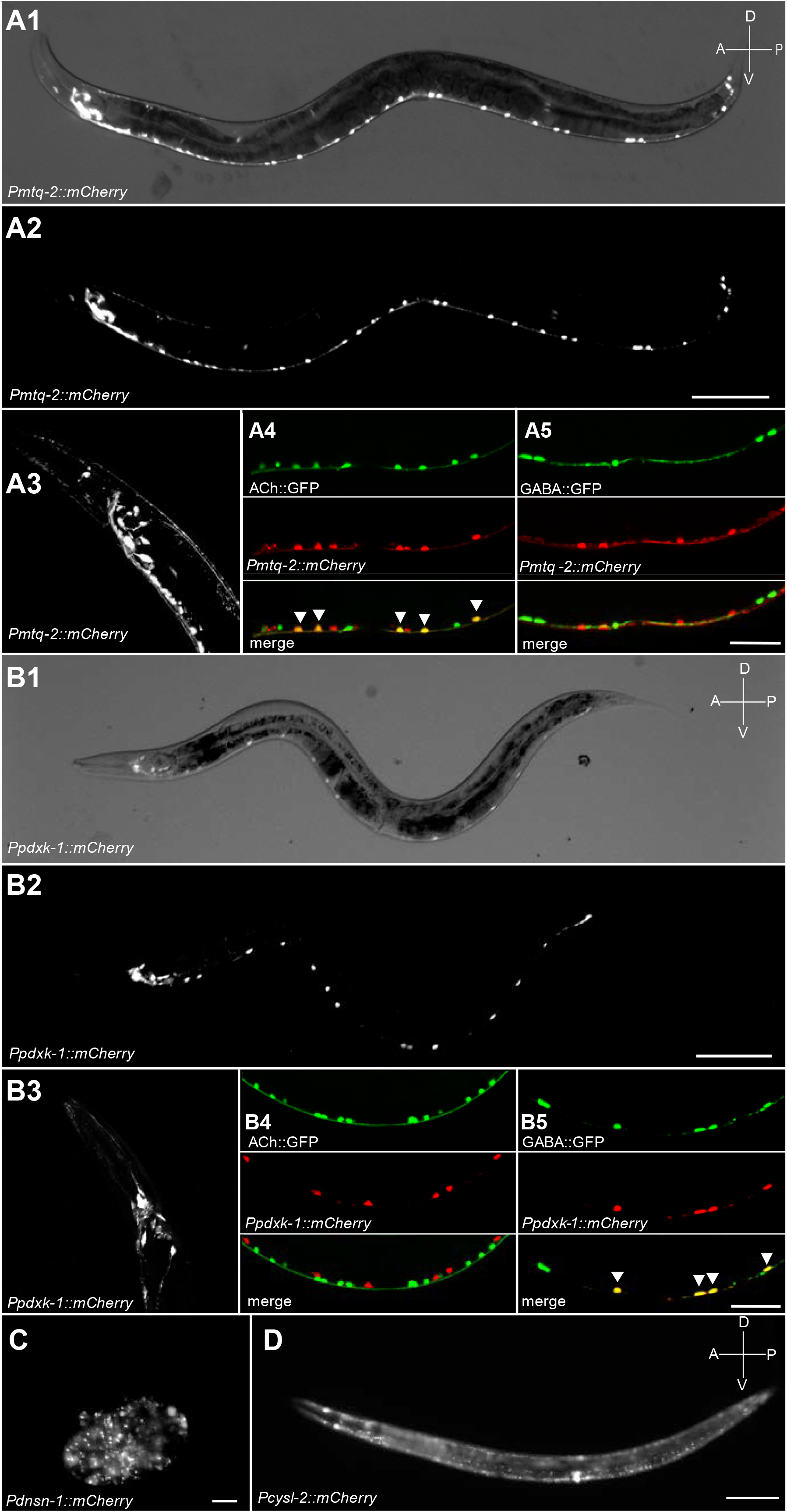
Expression pattern of select genes. Fluorescent micrographs of all HSA21 ortholog hits for which expression patterns were unknown. (**A1-3**) A transcriptional reporter for *mtq-2* expressed mCherry throughout the adult nervous system including in head (**A3**). Bright field image shown in **A1** for reference. Scale bar, 100μm. Dual color images show co-expression of *mtq-2* reporter in cholinergic (*arrowheads*, **A4**) but not in GABAergic neurons (**A5**) in the ventral nerve cord. Scale bar, 50 μm. (**B1-3**) A transcriptional reporter for *pdxk-1* expressed mCherry throughout the adult nervous system including in head (**B3**). Bright field image shown in **B1** for reference. Scale bar, 100 μm. Dual color images show co-expression of *pdxk-1* reporter in GABAergic neurons (**B5**) but not cholinergic neurons (**B4**) in the ventral nerve cord. Scale bar, 50 μm. (**C**) A transcriptional reporter for *dnsn-1* expressed mCherry widely in developing embryo. Scale bar, 50 μm. (**D**) A transcriptional reporter for *cysl-2* expressed mCherry throughout bodywall and vulval muscle. Scale bar, 100 μm.

**Table 1.**
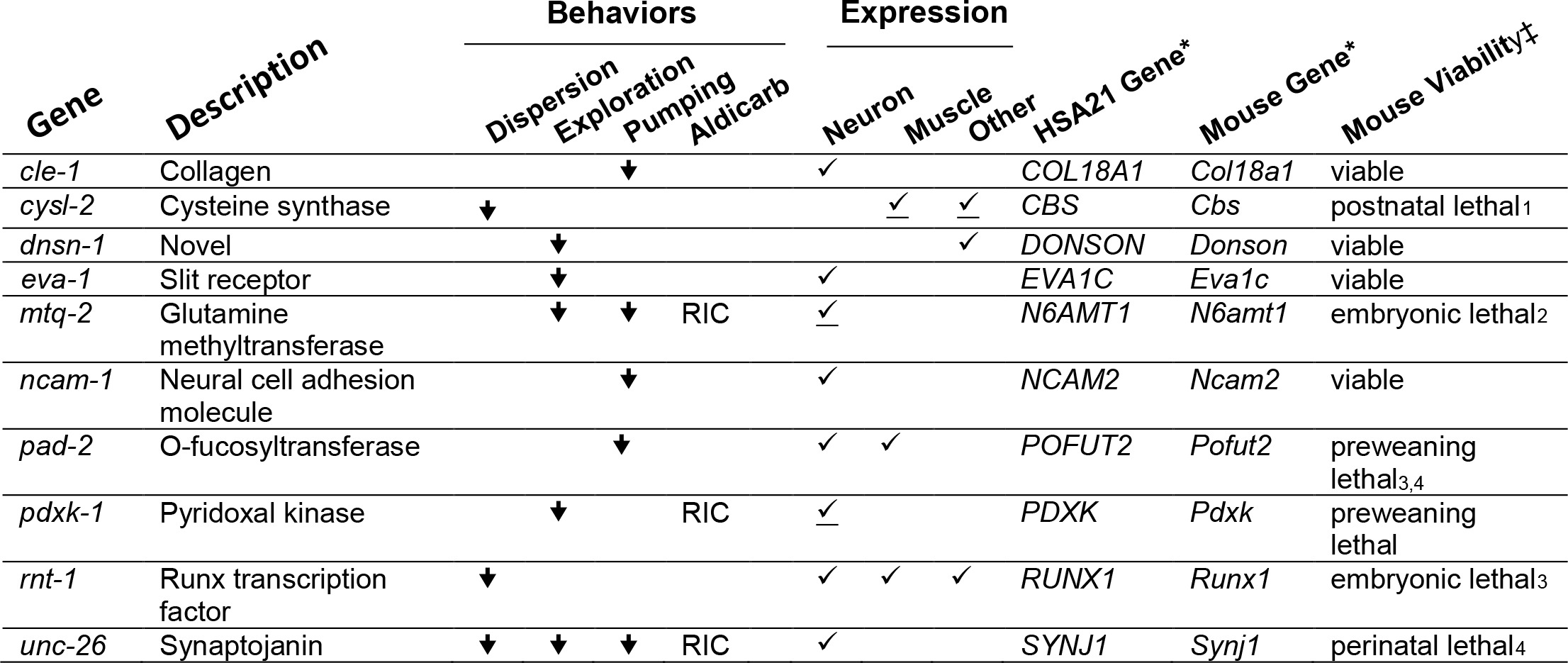
Summary of HSA21 gene orthologs required for normal behaviors in *C. elegans*. ↓ indicates the direction of changes that significantly differ from wild type (p < 0.001, one-way ANOVA) RIC (<u>R</u>esistant to <u>I</u>nhibitors of <u>C</u>holinesterase) indicates mutants with significant resistance to aldicarb *vs* wild type (p < 0.001, two-way ANOVA) ✓ indicates tissue(s) in which the gene is expressed. Data were derived from WormBase1. Underline signifies results found in this study. * HSA21 gene and mouse gene orthologs derived from InParanoid2. ‡ Mouse viability data derived from the Mouse Genome Informatics database7. Specific citations indicated with a footnote.

## DISCUSSION

Our research predicts new functional annotation of genes on the human 21^st^ chromosome by characterizing *in vivo* roles of putative orthologs in *C. elegans*. Within the field of Down syndrome, research has primarily focused on the functions of contiguous groups of genes using mice, such as those included in the so-called Down syndrome critical region (DSCR) (X. Jiang et al., 2015). Meanwhile, the vast majority of HSA21 genes remain functionally uncharacterized. One proven approach to reveal the function of large numbers of genes, even at the level of the whole genome, is to study their roles in more tractable invertebrate models such as *C. elegans*. Although several large-scale reverse genetics screens in *C. elegans* have begun to uncover novel *in vivo* roles for thousands of genes, they have focused less explicitly on neuronal phenotypes and excluded more genes than may be generally acknowledged (Fraser et al., 2000; Kamath et al., 2003; Sieburth et al., 2005; Vashlishan et al., 2008). For instance, two seminal whole-genome RNAi screens tested only 33 of the 51 HSA21 orthologs focused on in the current study (Fraser et al., 2000; Kamath et al., 2003). These studies primarily focused on viability phenotypes such as gross development and fertility. We expanded this analysis to test an additional eight orthologs not tested in these seminal or subsequent RNAi studies (Ceron et al., 2007; Gottschalk et al., 2005; Nakano et al., 2011; Pujol et al., 2001; Rual et al., 2004; Sönnichsen et al., 2005). With the addition of our current study, out of the 51 HSA21 orthologs, only five genes remain unstudied using RNAi or mutant in any study due to lack of reagents: *D1086.9 (CLIC6), H39E23.3 (CLIC6), Y54E10A. 11 (LTN1), Y74C10AL.2 (TMEM50B)*, and *ubc-14 (UBE2G2)*. Overall, however, our results were consistent with earlier studies. We found only one discrepancy, where a previous study found that RNAi targeting *irk-2* caused sterility while we found it had no effect (Kamath et al., 2003).

Other seminal RNAi screens in *C. elegans* searched for genes critical for synaptic transmission. Even though these screens were prodigious, testing 2,068 genes in total, they only tested five of the HSA21 orthologs: *mbk-1, itsn-1, ncam-1, K02C4.3*, and *wdr-4* (Sieburth et al., 2005; Vashlishan et al., 2008). These studies reported negative results for all five genes in agreement with our RNAi and mutant findings here. Notably, the synaptic RNAi screens did not test the three genes that we found to be critical for normal synaptic transmission: *mtq-2, pdxk-1*, and *unc-26*. Thus, despite the whole-genome and large-scale nature of previous screens, many genes with important *in vivo* functions likely remain to be discovered using *C. elegans*. Because our study largely replicated results from previous studies, however, we are optimistic that a systematic approach with *C. elegans*, as used here, may reveal the function of important sets of genes involved in human polygenic or chromosomal disorders.

We also performed several behavioral screens to identify HSA21 predicted orthologs that function in nervous system and muscle in ways that previous reverse-genetic screens may have missed. This included a radial dispersion assay that reflects short-term locomotor ability, an exploration assay that reflects longer-term sensory and motor integration, and a feeding assay that probes an independent neuromuscular circuit. We identified 10 mutants with deficiencies in these behaviors that we were able to rescue and/or confirm with independent alleles (**Table 1**): *cle-1 (COL18A1), cysl-2 (CBS), dnsn-1 (DONSON), eva-1 (EVA1C), mtq-2 (N6AMT1), ncam-1 (NCAM2), pad-2 (POFUT2), pdxk-1 (PDXK), rnt-1 (RUNX1)*, and *unc-26 (SYNJ)*. These 10 orthologs represent alluring candidates as genes important in nervous system development and/or function and should thus be prioritized for further study in worm and mammalian model systems.

In general, our targeted screen of HSA21 orthologs revealed three primary findings for the 10 HSA21 orthologs. First, we found genes with established links to nervous system function (e.g. *cle-1, eva-1, ncam-1*, and *unc-26)*. For instance, *cle-1 (COL18A1), eva-1 (EVA1C)*, and *ncam-1 (NCAM2)* and have all been linked to deficits in axon guidance in worm. The type XVIII collagen gene, *cle-1*, mediates both neuronal and axonal migrations (Ackley et al., 2001). *eva-1* encodes a Slit receptor required for axon guidance, and *ncam-1* encodes a neural cell adhesion molecule that functions redundantly with other cell adhesion molecules to direct axonal outgrowth (Fujisawa et al., 2007; Schwarz et al., 2009). Additionally, the *unc-26 (SYNJ)* mutant showed resistance to aldicarb but sensitivity to levamisole indicative of presynaptic defects. This is consistent with its known role in synaptic vesicle recycling (Harris et al., 2000).

Second, we uncovered novel neuronal functions for the otherwise well-characterized HSA21 ortholog *rnt-1 (RUNX1)*. In mammals, the runt-related transcription factor *RUNX1* plays a vital role in development, including regulation of the developing nervous system (Wang & Stifani, 2017). In worm, the runt-related transcription factor *rnt-1* has also been studied for its role in general development, including cell proliferation and male tail development (Hajduskova, Jindra, Herman, & Asahina, 2009; Nimmo, Antebi, & Woollard, 2005). Interestingly, loss of *rnt-1* may also cause defects to phasmid neuron morphology; however, this was not linked to a detectable behavioral phenotype (Hajduskova et al., 2009). The deficiency that the *rnt-1* mutant displayed in our radial dispersion assay might be due to developmental defects affecting the nervous system.

Third, whereas the majority of Down syndrome research has focused on well-studied genes, we also uncovered novel neuronal functions of three poorly characterized HSA21 orthologs that have weak or nonexistent links to the nervous system: *dnsn-1 (DONSON), mtq-2 (N6AMT1)* and *pdxk-1 (PDXK)*. These three genes are so poorly characterized that they only had placeholder names when we started our study. To encourage their study, we renamed them based on their predicted orthologs: *C24H12.5* to *dnsn-1, C33C12.9* to *mtq-2*, and *F57C9.1* to *pdxk-1*. Below, we highlight their functions in the limited context of their orthologs in other species.

*DONSON* was only recently found to encode a novel fork protection factor that underlies microcephalic dwarfism, yet it remains deeply understudied (Reynolds et al., 2017). The worm ortholog, *dnsn-1*, is equally uncharacterized. We found *dnsn-1* mutants to exhibit extended dwelling in an exploration assay and observed expression of *dnsn-1* in the developing embryo, but not the adult animal. Thus, our data are suggestive of a role within the nervous system and possibly within the developing nervous system.

To our knowledge, MTQ-2 has not been implicated in nervous system function in any animal model. The mammalian ortholog of *mtq-2, N6AMT1*, was originally named on the basis of the presence of an amino acid motif (D/N/S)PP(Y/FW) which is characteristic of <u>a</u>denine <u>m</u>ethyl<u>t</u>ransferases (Bujnicki & Radlinska, 1999; Kusevic, Kudithipudi, & Jeltsch, 2016). However, no evidence of adenine methylation by MTQ-2 protein has been found in either yeast or mouse, suggesting that *N6AMT1* may be a misnomer (Liu et al., 2010; Ratel et al., 2006). Instead, MTQ-2 has been shown to post-translationally modify ERF1 (<u>e</u>ukaryotic <u>r</u>elease <u>f</u>actor <u>1</u>) by methylating a universally conserved glutamine residue on ERF1 (Heurgué-Hamard et al., 2006); (Polevoda, Span, & Sherman, 2006). More recently, the mouse ortholog of MTQ-2 was shown to methylate many additional substrates *in vitro* and CHD5 (<u>c</u>hromodomain <u>h</u>elicase <u>D</u>NA-binding protein <u>5</u>) and NUT (<u>n</u>uclear protein in <u>t</u>estis) *in vivo*. This raises the possibility that MTQ-2 may modulate more proteins than previously thought (Kusevic et al., 2016). Here we have identified behavioral phenotypes of *mtq-2* mutants that support a neuronal role in *C. elegans*. Additionally, we found that a reporter for *mtq-2* widely expresses throughout the nervous system, specifically in a subset of cholinergic neurons. The high conservation (47% amino acid identity between human isoform 1 and worm) of MTQ-2 as well as many of its predicted interacting partners warrant more detailed study to determine how it functions in synaptic transmission. *C. elegans* may provide an especially informative system to study *mtq-2* as deletion of *N6AMT1* causes embryonic lethality in KO mice (Liu et al., 2010).

PDXK phosphorylates vitamin B6, converting it to PLP (pyridoxal-5’-phosphate), a key cofactor in the metabolism of hundreds of enzymatic reactions, including synthesis of neurotransmitters (Cao, Gong, Tang, Leung, & Jiang, 2006; Shetty & Gaitonde, 1980). Loss of *PDXK* in mice causes pre-weaning lethality, so its behavioral role has not yet been studied in mammals (Mouse Genome Informatics and the International Mouse Phenotyping Consortium (IMPC)). Interestingly, *PDXK* is predicted to be haploinsufficient in mammals and is a candidate dose-sensitive gene contributing to phenotypes in Down syndrome (Antonarakis, 2016). We found that *pdxk-1* mutant worms showed general deficiencies in behaviors as well as specific defects in synaptic transmission as determined by aldicarb screening. Our finding that the *pdxk-1* mutant was readily paralyzed by levamisole suggests a presynaptic role for *pdxk-1* in the nervous system.

There is a rich body of literature in the field of Down syndrome devoted to exploring the underlying genetic contributions of the disorder. Single gene studies have helped to elucidate the role of individual genes such as *DYRK1A*, both as a transgenic single gene triplication and within a trisomic context, and have led to the development of pharmacotherlapeutic treatments (Ahn et al., 2006; Altafaj et al., 2001; Shindoh et al., 1996; Song et al., 1996). Mouse models have also been instrumental in parsing discrete, chromosomal regions and uncovering genetic pathways that may be amenable to therapeutic targeting (P. V. Belichenko et al., 2015; Das et al., 2013; X. Jiang et al., 2015; Olson et al., 2004; 2007). More recently, bioinformatic approaches have been used to explore the transcriptional landscape of DS and highlight candidate genes that may contribute to DS phenotypes (Letourneau et al., 2014; Sturgeon, Le, Ahmed, & Gardiner, 2012). Our results provide initial clues on the function of several uncharacterized HSA21 orthologs that add to established findings. Conclusions on the precise mechanism of action of these genes and their possible relevance to DS will benefit from more detailed study in *C. elegans*—including examination of possible phenotypes induced by overexpression—as well as with further investigation using traditional mouse models and human cell line approaches.

## Supplemental information

Supplemental information found online

## Acknowledgements

We thank *Caenorhabditis* Genetic Center (funded by the NIH) and National Bioresource Project, Yishi Jin, Harald Hutter for strains. Susan Rozmiarek provided expert assistance, preliminary data were gathered by Allison Griffith, and Luisa Scott for constructive input. Funds provided by the Alzheimer’s Association, Down Syndrome Research and Treatment Foundation/LuMind, Research Down Syndrome, Point Rider Foundation, and NIH T-R01 Award 1R01AG041135 and 1RF1AG057355. Research was inspired by Ocean Pierce-Shimomura.

The authors declare no conflict of interest.

## Footnotes

1 Raymond Y N Lee et al., “WormBase 2017: Molting Into a New Stage.,” *Nucleic Acids Research*, October 24, 2017, doi:10.1093/nar/gkx998.

2 Erik L L Sonnhammer and Gabriel Östlund, “InParanoid 8: Orthology Analysis Between 273 Proteomes, Mostly Eukaryotic.,” *Nucleic Acids Research* 43, no. Database issue (January 2015): D234–39, doi:10.1093/nar/gku1203.

3 Judith A Blake et al., “Mouse Genome Database (MGD)-2017: Community Knowledge Resource for the Laboratory Mouse.,” *Nucleic Acids Research* 45, no. 1 (January 4, 2017): D723–29, doi:10.1093/nar/gkw1040.

1 M Watanabe et al., “Mice Deficient in Cystathionine Beta-Synthase: Animal Models for Mild and Severe Homocyst(E)Inemia.,” *Proceedings of the National Academy of Sciences* 92, no. 5 (February 28, 1995): 1585–89.

2 Peng Liu et al., “Deficiency in a Glutamine-Specific Methyltransferase for Release Factor Causes Mouse Embryonic Lethality.,” *Molecular and Cellular Biology* 30, no. 17 (September 2010): 4245–53, doi:10.1128/MCB.00218-10.

3 T Okuda et al., “AML1, the Target of Multiple Chromosomal Translocations in Human Leukemia, Is Essential for Normal Fetal Liver Hematopoiesis,” *Elsevier*

4 Ottavio Cremona et al., “Essential Role of Phosphoinositide Metabolism in Synaptic Vesicle Recycling,” *Cell* 99, no. 2 (October 1999): 179–88, doi:10.1016/S0092-8674(00)81649-9.

5 Raymond Y N Lee et al., “WormBase 2017: Molting Into a New Stage.,” *Nucleic Acids Research*, October 24, 2017, doi:10.1093/nar/gkx998.

6 Erik L L Sonnhammer and Gabriel Östlund, “InParanoid 8: Orthology Analysis Between 273 Proteomes, Mostly Eukaryotic.,” *Nucleic Acids Research* 43, no. Database issue (January 2015): D234–39, doi:10.1093/nar/gku1203.

7 Judith A Blake et al., “Mouse Genome Database (MGD)-2017: Community Knowledge Resource for the Laboratory Mouse.,” *Nucleic Acids Research* 45, no. 1 (January 4, 2017): D723–29, doi:10.1093/nar/gkw1040.

